# A spatially explicit delineation of transition zones for biogeographic regionalization

**DOI:** 10.64898/2025.12.16.693042

**Authors:** Rong Liu, Collin P. Gross, Barnabas H. Daru

**Affiliations:** Department of Biology, Stanford University, Stanford, CA, USA; Department of Invertebrate Zoology, National Museum of Natural History, Washington, DC, USA

**Author notes:** **Author of correspondence:** Rong Liu and Barnabas H. Daru. **Data availability:** All data and code required to reproduce the results have been uploaded to the journal’s submission system as supplementary files and available to editors and reviewers during peer review. **Authors’ Contributions**, R.L. conceived the study, analyzed the data and led the writing. B.H.D contributed to data analysis and writing. C.P.G. provided analytical feedback and contributed to writing. All authors approved the final manuscript.

**Keywords:** biogeography, biogeographic transition zones, transition zone index, hierarchical hard clustering, grade of membership model

## Abstract

1. A central goal of biogeography is to partition the world’s biota into meaningful biogeographical regions. While naturalists have long delineated biogeographic regions using hierarchical hard clustering approaches, they have often acknowledged the existence of transition zones. However, such transition zones have not been empirically identified or distinguished from hard clusters. Transition zones are outstanding areas for understanding the processes that shape biotic assembly; however, a coherent framework for delineating them is lacking.
2. Here, we propose a framework to empirically identify and analyze transition zones through a new metric, the “Transition Zone Index (TZI)”. The TZI decomposes the proportional contribution of multiple biotas to a spatial unit (called a map cell) derived from a Grade of Membership model into a single quantitative measure per cell using Shannon entropy principles. The transition zones can then be quantified and presented as a gradient, scaled from TZI = 0 (indicating distinctive bioregions) to TZI = 1 (corresponding to the highly transitional areas with maximally admixed biotas). Finally, we implement spatial regression models to detect the predictors of these transition zones.
3. We demonstrate the application of our framework using vascular plants of southern Africa. Through the spatial variation of TZI, our approach represents transition zones as a continuous gradient of biotic turnover between bioregions. These transition zones were found to be either previously delineated as discrete clusters or incorporated into neighboring clusters. Our approach effectively captures both the intensity and width of transitional dynamics, showing that these patterns are primarily shaped by precipitation.
4. Biogeographic transition zones are a measurable gradient, and our framework provides a robust quantitative foundation for detecting and analyzing the drivers of these areas. This improves our ability to understand the patterns and processes underlying biogeographic transition and provides important implications for conservation planning and biodiversity management.

## 1. Introduction

Biogeography is the study of the spatial and temporal distribution of biodiversity (Holt et al., 2013; Violle et al., 2014). A central goal of this field is to partition the world’s biota into meaningful biogeographic regions as the units for research in ecology, conservation and evolutionary biology (Jimenez et al., 2025; Liu et al., 2023). While early naturalists have devoted effort into identifying and delineating biogeographic regions since the 19th century, they have also acknowledged the existence of fuzzy transitional areas between biogeographic regions (Wallace, 1876). Contemporary biogeographic regionalization has predominantly relied on hierarchical clustering approaches which group biotas into geographic units bounded by hard boundaries (Jimenez et al., 2025; Liu et al., 2023). The drawback of these methods is the requirement that each spatial unit (sometimes called map cell) is allocated to a single region when in fact species assemblages in nature intergrade into each other (Kreft & Jetz, 2010). Despite the acknowledgement of transition zones in previous studies, they have not been empirically identified and distinguished from hard clusters.

Transition zones are commonly defined as the overlap areas between different biotas, which hold the characteristics of both regions and share a common evolutionary history (De Mendonça & Ebach, 2020; Morrone, 2024). These areas provide valuable opportunities for investigating biotic assembly mechanisms, as they contain a higher degree of biotic diversity and the complex patterns of intermixed biotas (Ferro & Morrone, 2014; Morrone, 2024). Researchers have developed multiple analytical approaches to identify transition zones: (i) Panbiogeographic track analysis (Urtubey et al., 2010) detects transition zones by the presence of intersections (nodes) or by conflicting results of the generalized distribution tracks derived from the congruent superposition of species’ individual tracks. (ii) Network community detection (Calatayud et al., 2019), applied to species-grid bipartite networks identifies modules (regions), with transition zones as the grid cells with low module specificity, that is, those with a low fraction of their links within assigned modules. (iii) Silhouette analysis (Jimenez et al., 2025) evaluates how securely each grid cell belongs to its cluster, and identifies the grid cells with low or negative silhouette values as uncertainty boundaries that indicate fuzzy borders. (iv) Species turnover methods (Mourelle & Ezcurra, 1997; Williams, 1996) identify transition zones by comparing each grid cell with its neighbors and highlight steep composition gradients, thereby revealing transition strength and width. Although the first three approaches can identify transition zones, they fail to maintain the inherently fuzzy nature of transition and end up assigning them to discrete clusters of their own or displaying them as crisp borders. The fourth approach, while able to measure the strength and width of transition zones, only quantifies transitional properties as between-cell dissimilarity rather than as a direct measure of biotic overlap that defines these important ecological interfaces.

To address these methodological gaps, we propose a framework to empirically delineate and analyze transition zones through a new metric, the “Transition Zone Index (TZI)”, which integrates outputs from Grade of Membership (GoM) model and Shannon entropy formula. The Grade of Membership model, originally devised for text-mining but now used in community ecology, can represent sampling units as having partial membership in multiple groups (Valle et al., 2014; White et al., 2019). It produces easily interpretable results, reflecting the underlying associations between species, branch lengths, or functional groups that admix to form local communities. This feature is important because it can allow the decomposition of assemblages into abrupt or gradual turnover along ecological gradients. This obviates the need to arbitrarily assign biomes to species, which has been the standard approach so far (Li et al., 2021; Valle et al., 2014; White et al., 2019). Although the GoM model identifies biotas, it insufficiently identifies transition zones themselves. Additionally, the lack of a standardized quantitative index to measure biotic intermixing constrains our ability to analyze spatial and temporal dynamics of transition zones, and to identify the abiotic and biotic factors that shape them.

In this study, we present our new TZI metric to empirically identify and analyze transition zones through GoM model into a single metric per cell using principles of Shannon entropy. Although Shannon entropy has been widely used in ecology to quantify community diversity by summarizing the species abundance distributions in a community (Chao et al., 2013; Sherwin & Prat i Fornells, 2019), we adapted its formula by incorporating the GoM-derived membership vectors instead of conventional species proportions. Therefore, TZI quantifies the compositional complexity of each map cell in terms of its biotic intermixing. A higher value indicates more complexity in biotic composition and thus a transitional state, while a lower TZI value reflects more distinctive regions dominated by single biotas. We apply the TZI framework to examine biogeographic patterns in vascular plant communities of southern Africa (Daru et al., 2015).

## 2. Materials and Methods

### 2.1 Study area and data

We used a presence-absence community matrix of vascular plants for southern Africa at a 50×50 km grid resolution. This dataset with 1,393 species in 365 map cells was from Daru et al. (2015), which has been included in R package *phyloregion* (Daru et al., 2020) as an example data in the package’s helpfile: ‘data(africa)’. This plant matrix M has entries *M*_*ji*_ ∈ {0,1}, where *j* indicates the map cells and *i* indicates species. A value of 1 means that species *i* is recorded in map cell *j*.

### 2.2 Analysis

#### (i) Grade of Membership model for soft regionalization

We inferred biogeographic regions using a Bernoulli GoM model (White et al., 2019). For each grid cell *j* and species *i*,

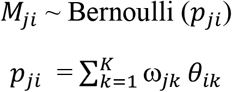

Here *p*_*ji*_ is the probability that vascular plant species *i* is present in the map cell *j*. The value of K is the number of the biotas fitted in the model. The membership vector ω_*j*_ = (ω_*j*1_, …, ω_*jK*_) gives the proportional contribution of each biota to map cell *j*, with ω_*jk*_ >= 0 and 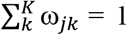. The parameter *θ*_*ik*_ ∈ [0,1] is the probability that species *i* occurs in the kth biota. This mixture of Bernoulli parameterization yields *p*_*ji*_∈ [0,1] and provides a low dimensional representation of community composition.

To determine the optimal number of biotas (*K)*, we fitted GoM models with different values of *K* ranging from 2 to 10, and the fit of each model was assessed in terms of Bayes factors such that higher Bayes Factors indicate an optimal *K*. The selected model with *K* = 4 achieved the best trade-off of predictive fit, stability and spatial coherence, and captures major biotic turnover of the southern Africa vascular plant assemblage (Fig. S1).

We used two different ways to map the biogeographic regions. First, we displayed the membership proportions vector ω_*j*_ as a pie chart centered on the centroid of the map cell *j* so that the proportional contribution of each biogeographic region can be visualized by the ratio of the representative color of each pie. However, this visualization introduces white interspaces between map cells because placing circular symbols on a regular grid leaves small unfilled areas between adjacent cells. Therefore, we improved the visualization by retaining square grid structure but representing each color with a blended color. This blended color was computed as a weighted combination of the base colors using the membership weights ω_*j*_. The analysis and visualization were performed in the R package *phyloregion* (Daru et al., 2020) using the function of *fitgom* to fit the GoM model, and visualized using function *plot_spatial_membership*, which by selecting parameter “type = ‘pie’” creates a pie chart map or “type = ‘blend’” which creates a blend color map.

#### (ii) Calculating the Transition Zone Index (TZI)

The Transition Zone Index provides a single metric for each map cell to quantify the degree of biotic overlap and thus identifying the transition zones along the biotic boundaries. This index adapts the Shannon entropy framework (Pielou, 1966) by substituting entropic component with membership vector derived from the GoM model in place of traditional species proportions.

For the map cell *j*, the membership vector ω_*j*_ = ( ω_*j*1_, …, ω_*jK*_ ) represents the proportional contribution across *K* biotas. The TZI is calculated as:

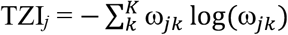

Following standard convention 0log0 = 0, thus ω_*jk*_= 0 contribute nothing to sum and requires no special handing. A high TZI indicates diffuse, multi-biota mixtures, while a low TZI indicates distinctive single-biota dominance.

To enable comparisons across biogeographic regionalization scheme with different number of biota *K*, we further normalized the TZI:

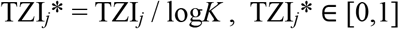

The normalized index TZI* = 1 occurs only when the contributions are perfectly even across *K* biotas, whereas TZI* = 0 indicates the contribution is entirely from a single biota (complete dominance). We visualized TZI* as a heatmap showing the degree of transition of each cell as a gradient from soft to hard regions to spatially delineate transition zones across southern Africa.

To directly estimate how many biotas effectively participate in each map cell, we also report the Hill number D (effective number of biotas):

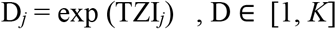

D = 1 indicates complete dominance by a single biota (corresponding to TZI* = 0), while D = K indicates the all *K* biotas contribute evenly (corresponding to TZI* = 1). This metric provides an intuitive interpretation of biotic complexity in units of equally abundant biotas.

To visualize the spatial distribution of transition zones at different levels of transitional degree, we mapped the biogeographic regions and the transition zones across TZI* larger than 0.65, 0.6. 0.55 and 0.5. The calculation of TZI has been included into the R package *phyloregion* as the function of *transition_zone_index(x)*, where *x* is omega, i.e., a matrix from the output of a grade of membership model.

#### (iii) Comparing hard clustering approach with soft clustering

We used hierarchical clustering as a hard-clustering baseline for vascular plants of southern Africa. The analyses were also performed in the R package *phyloregion* (Daru et al., 2020). First, we calculated the species-level beta diversity using the Simpson pairwise dissimilarity (Baselga, 2010) with the *beta_diss* function in *phyloregion*. To identify the most appropriate algorithms for our biogeographic regionalization schemes, we compared several commonly used hierarchical clustering algorithms, including UPGMA, UPGMC, WPGMC, WPGMA and others (Kreft & Jetz, 2010) using *select_linkage* function. We selected UPGMC as our clustering algorithm because it had the highest cophenetic coefficient, indicating that it preserved the best Simpson distance from the original distance matrix. We then used the elbow method with the function *optimal_phyloregion* to determine the optimal number of clusters based on the explained variance and internal consistency of clusters.

For comparison purposes, we set the number of UPGMC clusters *K* = 4 to match the GoM solution, as well as the optimal *K*= 13 of hierarchical clustering itself. To visualize differences between soft and hard clustering approaches and highlight transition zones, we overlaid the outlines identified by hard clustering onto the TZI heatmap.

#### (iv) Identifying the predictors shaping transition zones

We included two types of predictor variables, derived from the WorldClim database (Fick & Hijmans, 2017). The first type is the climate conditions, including mean annual temperature, temperature seasonality, mean annual precipitation, precipitation seasonality and others. The second type is topographic conditions, including elevation, terrain ruggedness index (TRI), terrain position index (TPI) and others. The full list of all predictors is provided in Table S1. All predictor variables were z-standardized (mean = 0, SD = 1) before analysis.

To address potential multicollinearity among predictor variables, we calculated the variance inflation factors (VIF) to retain only independent variables which VIF were less than ten. We then examined the predictors of transition zones while accounting for spatial dependence by fitting the spatial regression models to the selected predictor variables with normalized index TZI*using the R package *spatialreg* (Bivand et al., 2021). We built a k-nearest neighbors (k = 8) spatial weights structures to ensure each map cell was connected to its eight nearest neighbors. We compared four models: spatial lag, spatial error, spatial Durbin, and spatial autoregressive combined models. Model selection based on Akaike information criterion (AIC) showed that the spatial autoregressive combined model was optimal. Then we additionally computed average direct, indirect and total impacts using *impacts* function. The average total impacts from the selected model were reported as the standardized effect sizes as all the predictor variables were standardized.

## 3. Results

The GoM model with *K* = 4 achieved the best trade-off of predictive fit, stability and spatial coherence, and effectively captures major biotic turnover pattern of plant communities of southern Africa (Fig. S1). The largest biogeographic region corresponded closely to the distribution of Savanna-Namib Desert, followed by Miombo Woodlands, Fynbos and Grassland (Fig. 1). These biogeographic region designations follow the regionalization scheme of Daru et al. (2016).

**Fig. 1.**
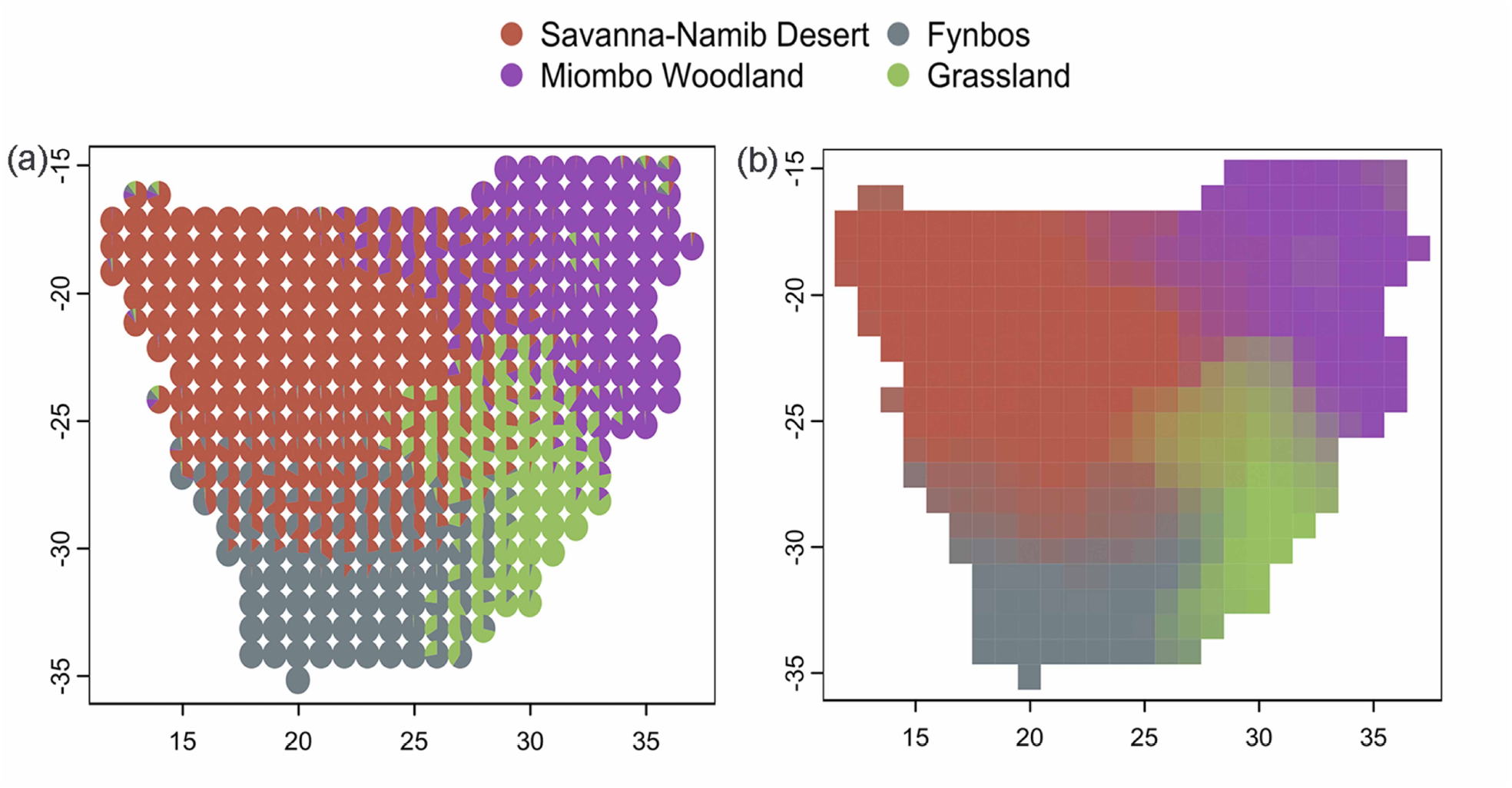
Biogeographic regionalization of plant species across southern Africa using Grade of Membership model. (a) Pie chats represent the proportional contribution of biotas to each 50 × 50 km grid cell. (b) Same as (a) but using blended colors to show continuous turnover between biotas, obviating the need for pie charts. Number of biotas *K* = 4. Different colors present different biotas.

The TZI heatmap shows the degree of transition of each map cell as a gradient ranging from transitional (red) to distinctive (blue) regions (Fig. 2). The proportional contribution of the dominant biota to each map cell decreased as the TZI increased (Fig. S3). No area shows complete mixing by all four biotas, with the largest intermixing comprising three biotas with a TZI of 0.79 (and contributions of 0.26, 0.33 and 0.40, for each biota, respectively). The lowest TZI of 0.017 was characterized by a dominant biota with a high proportional contribution of 0.997.

**Fig. 2.**
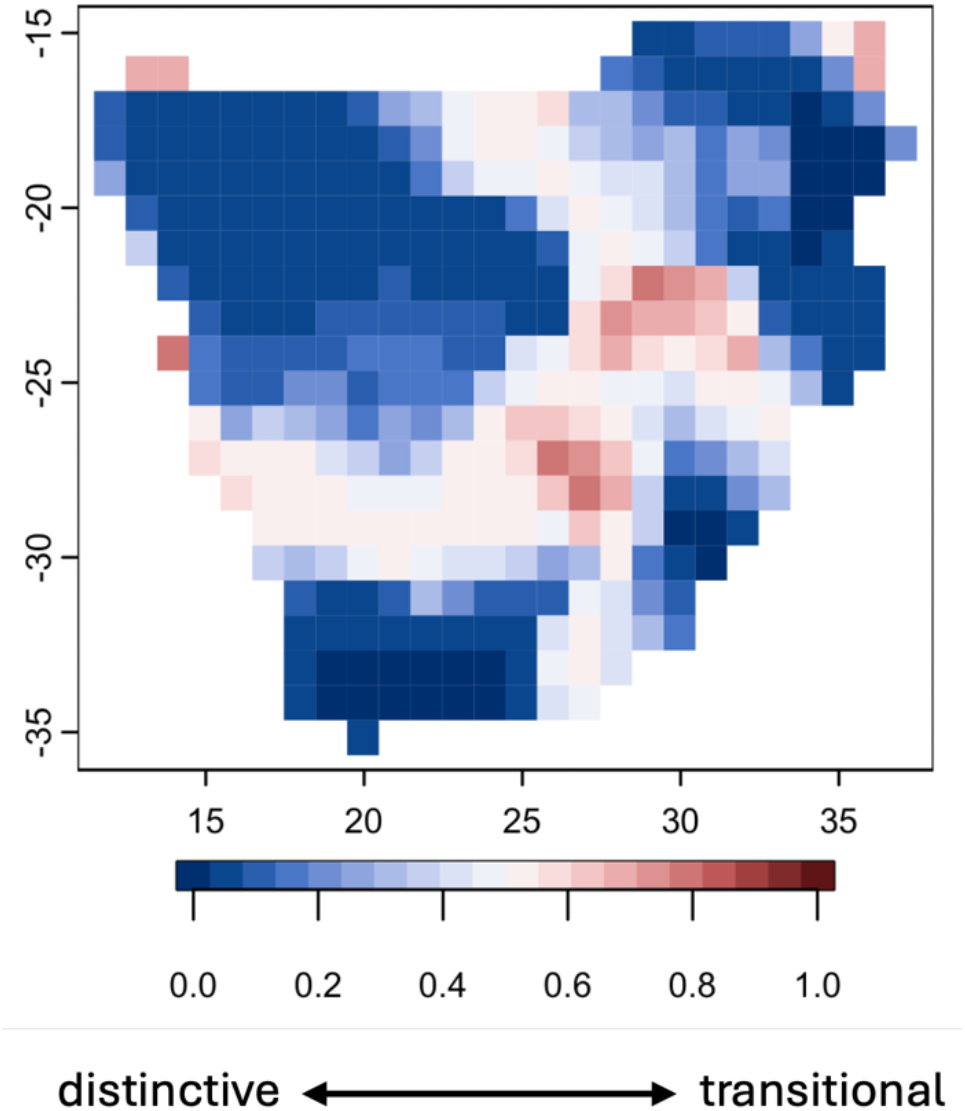
Transition Zone Index (TZI) showing the strength of transition zones as a gradient from distinctive to transitional areas. The normalized TZI shows a gradient, scaled from 0 indicating distinct biogeographic regions to 1 corresponding to transition zones where biotas admixed.

The spatial distribution map shows how the intensity and width of transition zones change across different TZI threshold levels (Fig. 3). The strongest transitions (TZI > 0.6) occur in a few convergence areas of three biotas, forming discrete transitional hotspots. As the TZI threshold decreases (0.55 to 0.50), the transition zones become interconnected and form continuous belts. The width of transition zones was also found to be broader in convergence areas, but narrower along the biotic boundaries between two biotas.

**Fig. 3.**
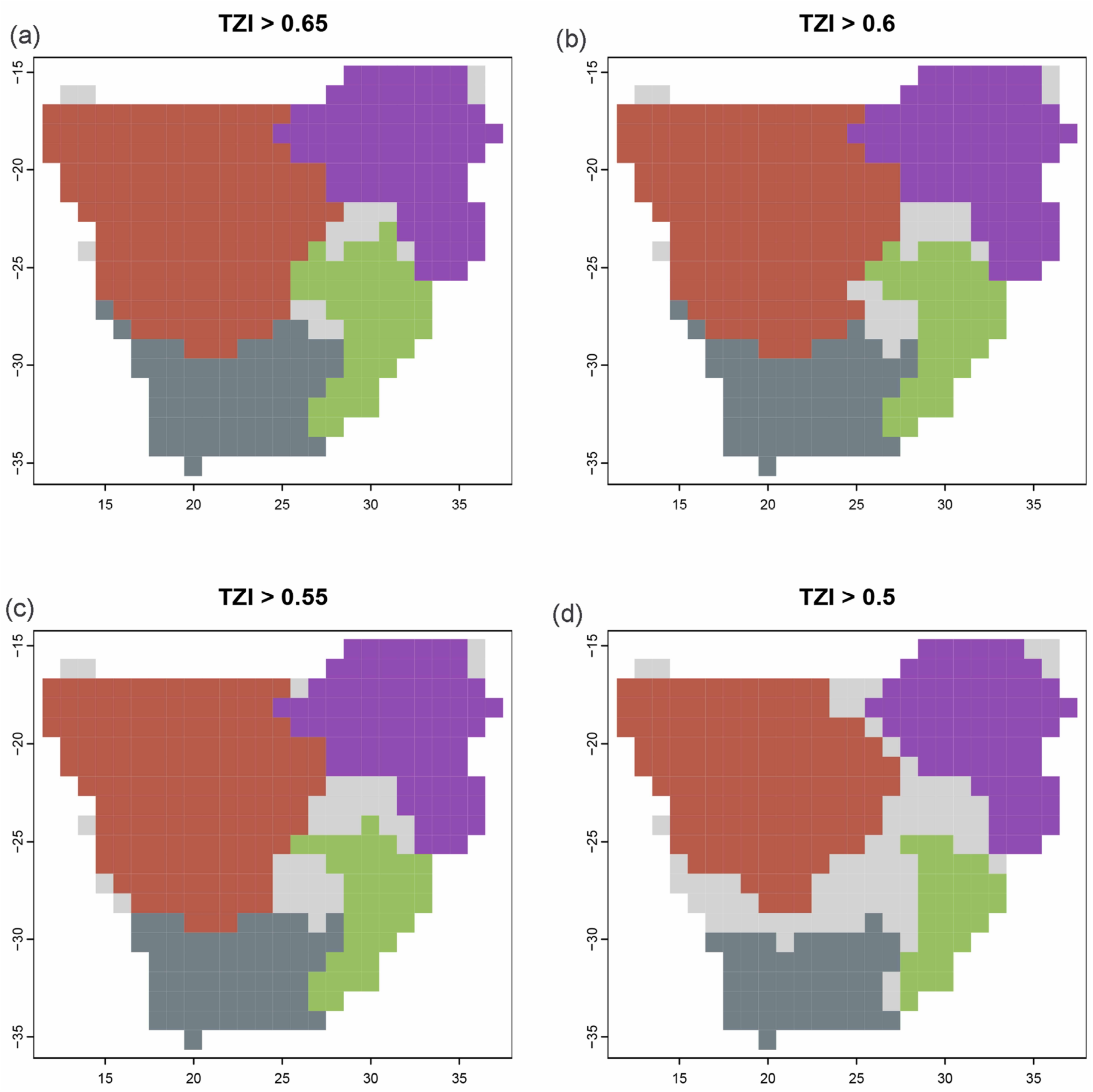
Spatial distribution of transition zones across varying Transition Zone Index (TZI). (a-d) Light grey indicates map cells with TZI > 0.65, 0.6, 0.55, 0.5. The colored polygons represent dominant biotas, and the color of each biota is the same as in Fig 1. Map cells with an TZI greater than certain thresholds are assigned a light grey color, whereas cells with TZI below thresholds are depicted in the color representing the biotas making the largest proportional contribution.

On the other hand, hierarchical clustering and the elbow method revealed that a division of 13 clusters explained 76% of the variation in beta diversity across map cells (Fig. S2). The overlay of the outlines of the hard clusters onto the TZI heatmap revealed that some areas which had been identified as distinctive regions were in fact transition zones. For example, the transition zones between the Savanna-Namib Desert and Fynbos appeared as discrete clusters in hierarchical clustering. Meanwhile some transition zones had been forcibly incorporated into one of the neighboring hard clusters. For example, the transition zone between Savanna-Namib Desert and Miombo Woodlands was included as part of the Savanna-Namib Desert (Fig. 4).

**Fig. 4.**
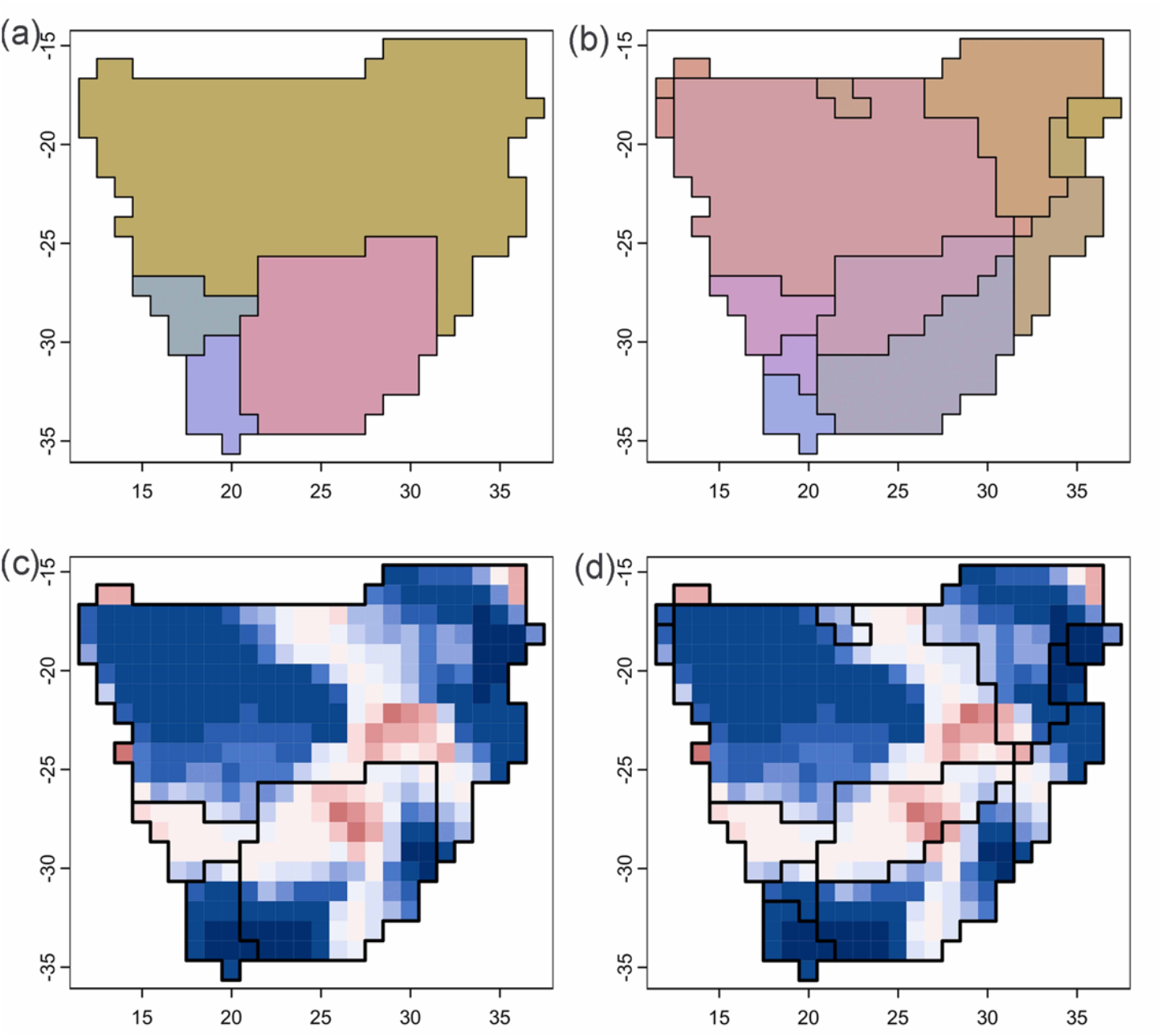
Comparing the hard clustering approach with the Grade of Membership (GoM) model. (a) Traditional hard clustering based on a hierarchical cluster analysis of a beta diversity matrix, with *K* set to 4 to match the GoM approach. (b) Same as (a), but with K = 13 (the optimal clusters for which hierarchical clustering preserves 76% of the variation in beta dissimilarity. (c-d) Overlay of the outlines of the hard clusters onto the GoM approach reveals that some areas delineated by the hard clustering approach as the discrete clusters are in fact transitions zones.

The best-fit model being spatial autocorrelation combined, successfully controlled the spatial autocorrelation of TZI. The results show that precipitation rather than temperature or topography, plays a significant role in shaping transition zones in southern Africa (Fig. 5). The areas where precipitation was more stable had lower TZI.

**Fig. 5.**
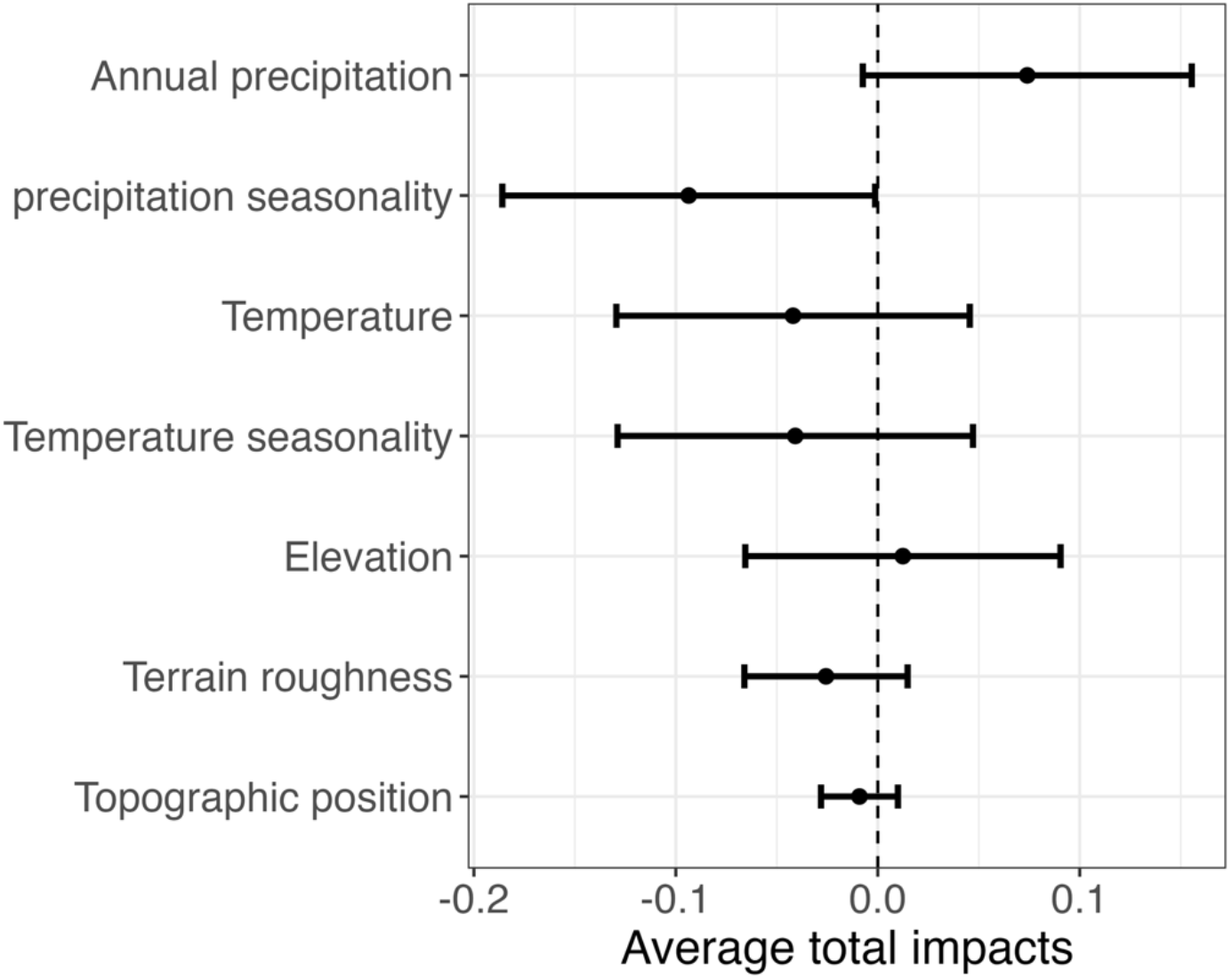
Effects of environmental variables on the Transition Zone Index (TZI) based on spatial autoregressive combined model. The average total impacts (standardized units) indicate the relative effect of each predictor on the TZI.

## 4. Discussion

Biogeographic transition zones are outstanding areas for investigating the evolutionary processes that shape biotic assembly (Morrone, 2024). However, there has been no coherent framework for studying transition zones until now. We propose a new framework which identifies and analyzes biogeographic transition zones based on the GoM model and Shannon entropy. Our framework depicts a more realistic and natural representation of transition zones, as continuous gradients of biotic turnover between bioregions. Through the spatial variation of TZI, our approach effectively captures transitional dynamics in terms of both intensity and spatial extent.

Our framework has several advantages. First, it portrays a more realistic and natural transition of biogeographic regions as a continuous gradient of biotic turnover between regions. The classification of a transition zone has varied considerably over time (Morrone, 2024), with transitions zones treated either as a subregion and assigned to one of the regions involved (Wallace, 1876), considered as separate discrete regions (De Mendonça & Ebach, 2020), not assigned to any of the regions (Simpson, 1977); or assigned to both regions simultaneously (Morrone, 2015). In our study, rather than engaging with the complex debate surrounding transition zone classification, we avoided assigning the transition zones to any of these classes. We instead depicted the continuous spatial turnover pattern of transition zones as a gradient from distinctive to transitional areas. We used the Transition Zone Index (TZI) to present this gradient as a scale from 0 indicating distinct biogeographic regions to 1 corresponding to transition zones where biotas are admixed. The transition zones we identified in southern Africa with the TZI are consistent with previous consensus on vegetation structure and composition in southern Africa, which identified increasing plant community structural complexity and species changes from the Northern Cape Province of South Africa to the northeast of Botswana and the Caprivi area of Namibia (Scholes et al., 2002). This continuous depiction of transition zones overcomes the limitations of traditional hard clustering approach, which identifies transition zones as distinctive regions or forcibly incorporates them into one of the neighboring clusters (Fig. 4).

Second, our approach effectively captures transitional dynamics in terms of both intensity and spatial extent through the spatial variation of TZI. Previous approaches have either assigned transition zones as discrete regions (e.g., using track analysis or network modules) (Urtubey et al., 2010; Calatayud et al., 2019) or portrayed them as crisp borders (silhouette analysis) (Jimenez et al., 2025) thus failing to measure the intensity and the width of transition zones. The species turnover approach has been the only method that can estimate strength and width of transition zones, but it operates on dissimilarity between neighboring map cells rather than directly quantifying biotic overlap (Mourelle & Ezcurra, 1997; Williams, 1996). By contrast, TZI is computed from the proportional contribution of multiple biotas to each cell, therefore fitting with the definition of transition zones as areas of overlap (De Mendonça & Ebach, 2020; Morrone, 2024). In southern Africa, although our framework identified four distinctive bioregions, the strongest transitions occurred in a few specific convergence areas (Fig. 2). These transitions involve the intermixing of only three of the four biogeographic regions, resulting in the formation of two discrete transitional hotspots. The two hotspots were separated by a narrow corridor comprising biotas from the Savanna-Namib Desert and Grassland biomes. Beyond the transitional intensity, our approach can also identify both abrupt and gradual transition zones between these bioregions. The broadest and most gradual transition occurred in the same area as transitional hotspots, followed by a connected gradual tunnel between the Savanna-Namib Desert and Fynbos, while the transition zone between the Savanna-Namib Desert and Miombo Woodlands was also extensively connected but narrower. The transition zones between the Miombo Woodlands and Grassland, and between the Fynbos and Grassland are short, abrupt and unconnected.

Thirdly, our TZI provides a quantitative method for detecting the drivers of transition zones by providing a single metric for each map cell. Due to the lack of a single metric to represent the degree of transition, previous analyses focused on the drivers that influence the presence or absence of biogeographic boundaries (Ficetola et al., 2017; Gross et al., 2025; Minev-Benzecry & Daru, 2024), which involves using a binary variable to determine whether a given cell is in contact with a biogeographic region boundary or not as the response variable and then fitting a generalized model to test the effects of the drivers. Our TZI approach alternatively, identifies the drivers which cause an area to be more transitional. Tectonics and climate (especially climate seasonality) are primary factors determining biogeographic boundaries, with the former typically creating deep boundaries between biogeographic realms across continents, while the latter more often influencing shallow boundaries within continents (Antonelli, 2017; Ficetola et al., 2017; Liu et al., 2023). Our results further demonstrated that climate seasonality, specifically precipitation seasonality was the major predictor driving intensity of transition zones in southern Africa (Fig. 5). This finding is reasonable, as southern Africa comprises vast arid and semiarid areas, where the drought and floods caused by precipitation fluctuation impose significant pressure on plant communities (Geppert et al., 2022; Scholes et al., 2002). Consequently, plants tend to distribute in areas with less variable precipitation thus developing the transition zones.

Together, our framework represents biogeographic transition zones as measurable gradients, overcoming the major limitations of categorical approaches by quantifying the intensity and width of transition zones and links these patterns to environmental drivers. Although our demonstration of the TZI framework is with plants of southern Africa, this methodology can be extended to different organismal groups, for example, animal and microbiome communities, and across different geographic scales. The normalized index TZI* eliminates the effect of the number of biotas (K), which enables standardized comparisons across biomes, and provides a unified quantitative basis for evaluating similarities and differences in transition zones among geographic regions and taxonomic groups. While we only presented contemporary spatial patterns of transition zones, the framework can also be applied to track temporal dynamics of transition zones through reconstructions of historical species distributions. Importantly, rather than tracking the changes in each individual biota, we can directly focus on the contraction, expansion, or movement of transition zones themselves in a way that can capture the spatial and temporal changes of different biotas. Overall, our TZI framework provides a quantitative perspective for studying biogeographic transition zones that improves our ability to understand the spatial patterns of biodiversity and their dynamic changes. Future work should explore the transition zones across different spatial scales, and their association with the species evolutionary history and ecological functionality.

**Table S1.** The climate and topographic predictors used in this study.

| Climate predictors | Description |
| --- | --- |
| BIO1 = Annual Mean Temperature |  |
| BIO2 = Mean Diurnal Range | Mean of monthly (max temp - min temp) |
| BIO3 = Isothermality | $(\text{BIO2}/\text{BIO7}) \times 100$ |
| BIO4 = Temperature Seasonality | standard deviation $\times 100$ |
| BIO5 = Max Temperature of Warmest Month |  |
| BIO6 = Min Temperature of Coldest Month |  |
| BIO7 = Temperature Annual Range | BIO5-BIO6 |
| BIO8 = Mean Temperature of Wettest Quarter |  |
| BIO9 = Mean Temperature of Driest Quarter |  |
| BIO10 = Mean Temperature of Warmest Quarter |  |
| BIO11 = Mean Temperature of Coldest Quarter |  |
| BIO12 = Annual Precipitation |  |
| BIO13 = Precipitation of Wettest Month |  |
| BIO14 = Precipitation of Driest Month |  |
| BIO15 = Precipitation Seasonality | Coefficient of Variation |
| BIO16 = Precipitation of Wettest Quarter |  |
| BIO17 = Precipitation of Driest Quarter |  |
| BIO18 = Precipitation of Warmest Quarter |  |
| BIO19 = Precipitation of Coldest Quarter |  |
| Topographic predictors | Description |
| Elevation |  |
| Slope |  |
| Terrain ruggedness index (TRI) | Mean absolute elevation difference between a focal cell and its eight neighboring cells |
| Terrain position index (TPI) | Elevation of a cell minus the mean elevation of its eight neighboring cells (positive = ridges; 0 = flat; negative = valleys) |
| Roughness | Elevation range with the neighboring cells |

**Fig. S1.**
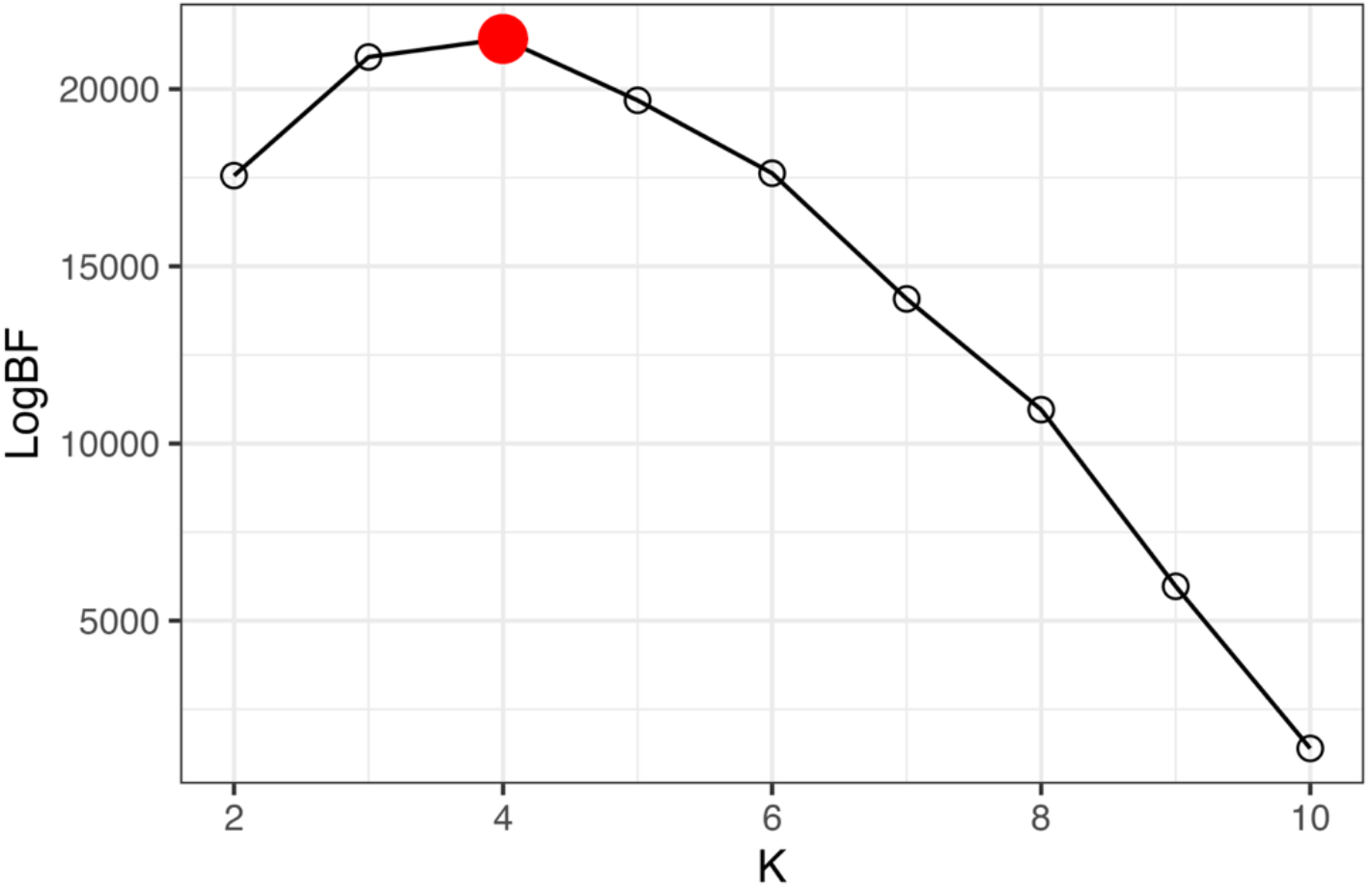
Grade of Membership model to determines the optimal number of biotas based on the Bayes Factor for a varying number of K. The optimal number of biotas is 4, indicated in red.

**Fig. S2.**
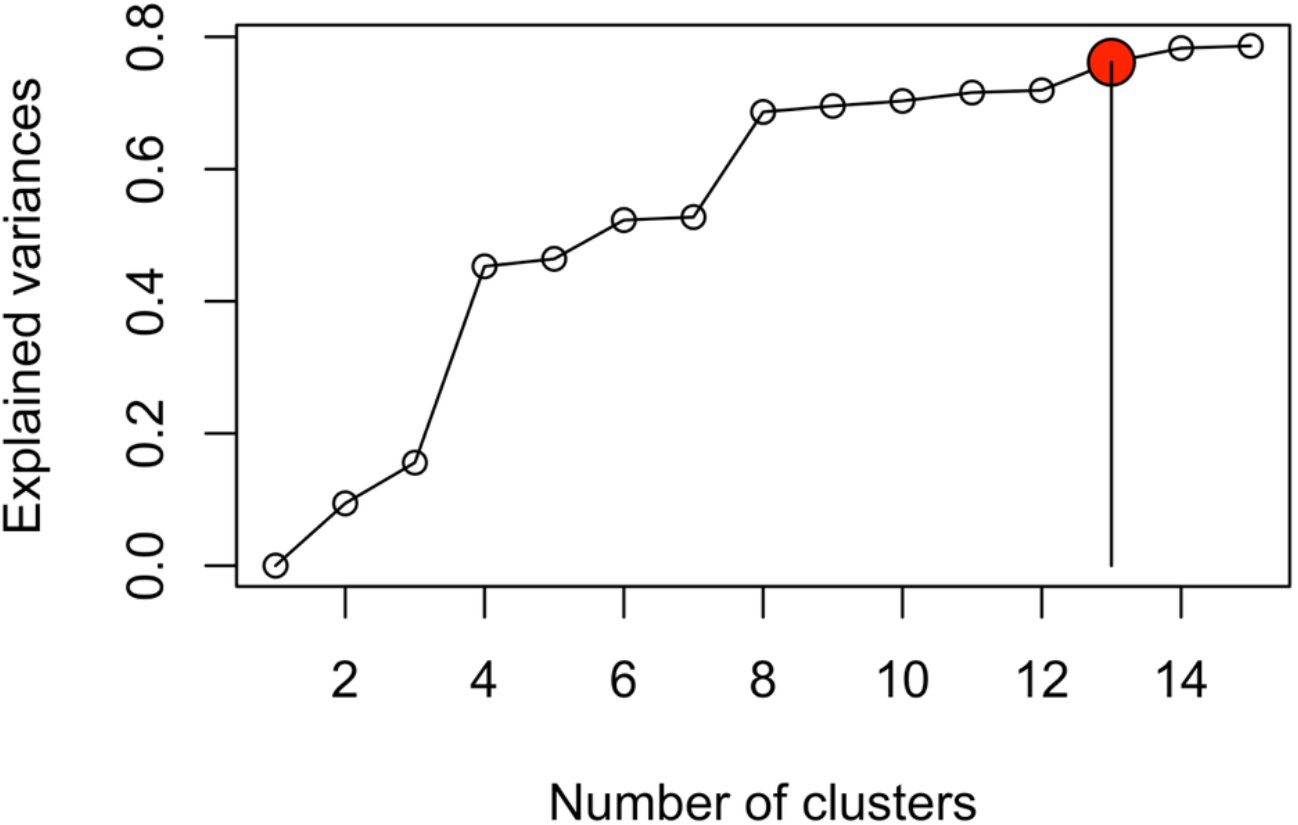
Hierarchical hard clustering approach to determine the optimal number of clusters based on the elbow method. The optimal number of clusters is 13, indicated in red.

**Fig. S3.**
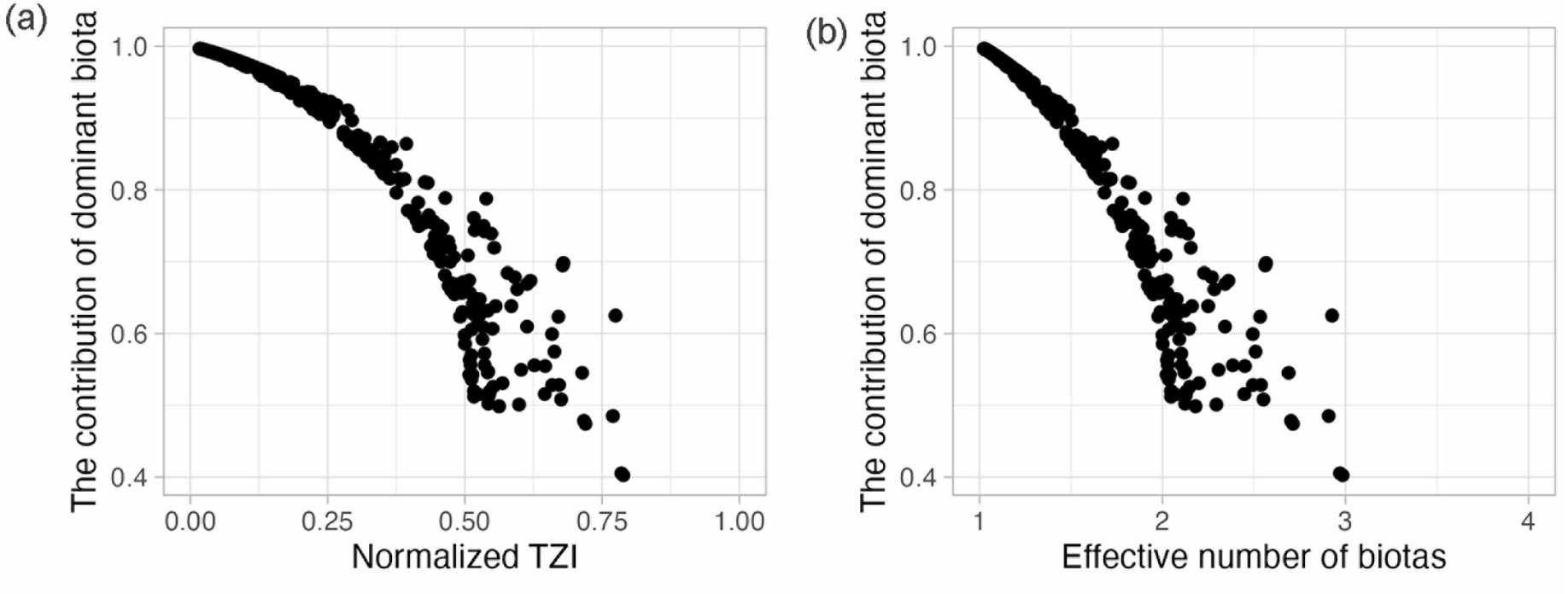
The relationship between the contribution of dominant biotas to each map cell and normalized TZI (a) and effective number of biotas (b).

## Acknowledgement

We thank Xianyu Yang, Joseph Kesler, Lauren Puleo, and Paul Markley for helpful comments and suggestions. This project was supported by grants from the U.S. National Science Foundation (awards 2345994 and 2416314), Alfred P. Sloan Foundation, and Stanford Woods Institute Big Ideas for Oceans.

## Notes

### Competing Interest Statement

The authors have declared no competing interest.

## References

Antonelli, A. (2017). Biogeography: Drivers of bioregionalization. Nature Ecology & Evolution, 1(4), 0114. 10.1038/s41559-017-0114

Bivand, R., Millo, G., & Piras, G. (2021). A review of software for spatial econometrics in R. Mathematics, 9(11), 1276. 10.3390/math9111276

Calatayud, J., Rodríguez, M.Á., Molina-Venegas, R., Leo, M., Horreo, J. L., & Hortal, J. (2019). Pleistocene climate change and the formation of regional species pools. Proceedings of the Royal Society B: Biological Sciences, 286(1905), 20190291. 10.1098/rspb.2019.0291

Chao, A., Wang, Y. T., & Jost, L. (2013). Entropy and the species accumulation curve: A novel entropy estimator via discovery rates of new species. Methods in Ecology and Evolution, 4, 1091–1100. 10.1111/2041-210X.12108

Daru, B. H., Karunarathne, P., & Schliep, K. (2020). phyloregion: R package for biogeographical regionalization and macroecology. Methods in Ecology and Evolution, 11(11), 1483– 1491. 10.1111/2041-210X.13478

Daru, B. H., van der Bank, M., & Davies, T. J. (2015). Spatial incongruence among hotspots and complementary areas of tree diversity in southern Africa. Diversity and Distributions, 21(7), 769–780. 10.1111/ddi.12290

Daru, B. H., van der Bank, M., Maurin, O., Yessoufou, K., Schaefer, H., Slingsby, J. A., & Davies, T. J. (2016). A novel phylogenetic regionalization of phytogeographical zones of southern Africa reveals their hidden evolutionary affinities. Journal of Biogeography, 43(1), 155–166. 10.1111/jbi.12619

Ferro, I., & Morrone, J. J. (2014). Biogeographical transition zones: A search for conceptual synthesis. Biological Journal of the Linnean Society, 113(1), 1–12. 10.1111/bij.12333

Ficetola, G. F., Mazel, F., & Thuiller, W. (2017). Global determinants of zoogeographical boundaries. Nature Ecology & Evolution, 1(4), 0089. 10.1038/s41559-017-0089

Fick, S. E., & Hijmans, R. J. (2017). WorldClim 2: New 1-km spatial resolution climate surfaces for global land areas. International Journal of Climatology, 37(12), 4302–4315. 10.1002/joc.5086

Geppert, M., Hartmann, K., Kirchner, I., Pfahl, S., Struck, U., & Riedel, F. (2022). Precipitation over southern Africa: Moisture sources and isotopic composition. Journal of Geophysical Research: Atmospheres, 127(21), e2022JD037005. 10.1029/2022JD037005

Gross, C. P., Wright, A. M., & Daru, B. H. (2025). A global biogeographic regionalization for butterflies. Philosophical Transactions of the Royal Society B: Biological Sciences, 380(1917), 20230211. 10.1098/rstb.2023.0211

Hermogenes De Mendonça, L., & Ebach, M. C. (2020). A review of transition zones in biogeographical classification. Biological Journal of the Linnean Society, 131(4), 717– 736. 10.1093/biolinnean/blaa120

Holt, B. G., Lessard, J.-P., Borregaard, M. K., Fritz, S. A., Araújo, M. B., Dimitrov, D., Fabre, P.-H., Graham, C. H., Graves, G. R., Jønsson, K. A., Nogués-Bravo, D., Wang, Z., Whittaker, R. J., Fjeldså, J., & Rahbek, C. (2013). An update of wallace’s zoogeographic regions of the world. Science, 339(6115), 74–78. 10.1126/science.1228282

Jimenez, Y. G., Paz, A., Bialic-Murphy, L., Crowther, T., & Maynard, D. S. (2025). A Global Regionalisation of Tree Functional Capacity. Global Ecology and Biogeography, 34(7), e70083. 10.1111/geb.70083

Kreft, H., & Jetz, W. (2010). A framework for delineating biogeographical regions based on species distributions. Journal of Biogeography, 37(11), 2029–2053. 10.1111/j.1365-2699.2010.02375.x

Li, Q., Sun, H., Boufford, D. E., Bartholomew, B., Fritsch, P. W., Chen, J., Deng, T., & Ree, R. H. (2021). Grade of Membership models reveal geographical and environmental correlates of floristic structure in a temperate biodiversity hotspot. New Phytologist, 232(3), 1424–1435. 10.1111/nph.17443

Liu, Y., Xu, X., Dimitrov, D., Loic Pellissier, Borregaard, M. K., Shrestha, N., Su, X., Luo, A., Zimmermann, N. E., Rahbek, C., & Wang, Z. (2023). An updated floristic map of the world. Nature Communications, 14(1), 2990. 10.1038/s41467-023-38375-y

Minev-Benzecry, S., & Daru, B. H. (2024). Climate change alters the future of natural floristic regions of deep evolutionary origins. Nature Communications, 15(1), 9474. 10.1038/s41467-024-53860-8

Morrone, J. J. (2015). Biogeographical regionalisation of the world: A reappraisal. Australian Systematic Botany, 28(3), 81–90. 10.1071/SB14042

Morrone, J. J. (2024). Why biogeographical transition zones matter. Journal of Biogeography, 51(4), 544–549. 10.1111/jbi.14632

Mourelle, C., & Ezcurra, E. (1997). Differentiation diversity of argentine cacti and its relationship to environmental factors. Journal of Vegetation Science, 8(4), 547–558. 10.2307/3237206

Pielou, E. C. (1966). The measurement of diversity in different types of biological collections. Journal of Theoretical Biology, 13, 131–144. 10.1016/0022-5193(66)90013-0

Scholes, R. j., Dowty, P. r., Caylor, K., Parsons, D. a. b., Frost, P. g. h., & Shugart, H. h. (2002). Trends in savanna structure and composition along an aridity gradient in the kalahari. Journal of Vegetation Science, 13(3), 419–428. 10.1111/j.1654-1103.2002.tb02066.x

Sherwin, W. B., & Prat i Fornells, N. (2019). The introduction of entropy and information methods to ecology by ramon margalef. Entropy, 21(8), 794. 10.3390/e21080794

Simpson, G. G. (1977). Too many lines; the limits of the oriental and australian zoogeographic regions. Proceedings of the American Philosophical Society, 121(2), 107–120.

Urtubey, Estrella, Stuessy, T., Tremetsberger, K., & Morrone, J. (2010). The south american biogeographic transition zone: An analysis from asteraceae. Taxon, 59, 505–509. 10.2307/25677608

Valle, D., Baiser, B., Woodall, C. W., & Chazdon, R. (2014). Decomposing biodiversity data using the latent dirichlet allocation model, a probabilistic multivariate statistical method. Ecology Letters, 17(12), 1591–1601. 10.1111/ele.12380

Violle, C., Reich, P. B., Pacala, S. W., Enquist, B. J., & Kattge, J. (2014). The emergence and promise of functional biogeography. Proceedings of the National Academy of Sciences, 111(38), 13690–13696. 10.1073/pnas.1415442111

White, A. E., Dey, K. K., Mohan, D., Stephens, M., & Price, T. D. (2019). Regional influences on community structure across the tropical-temperate divide. Nature Communications, 10(1), 2646. 10.1038/s41467-019-10253-6

Williams, P. H. (1996). Mapping variations in the strength and breadth of biogeographic transition zones using species turnover. Proceedings: Biological Sciences, 263(1370), 579–588.

Wallace, A. R. (1876). The geographical distribution of animals; with a study of the relations of living and extinct faunas as elucidating the past changes of the Earth’s surface. New York: Harper & Brothers. Volume 1.

